# Calcimycin mediates apoptosis in Breast and Cervical cancer cells by inducing intracellular calcium levels in a P2RX4-dependent manner

**DOI:** 10.1101/2023.07.06.548052

**Authors:** Neha, Prashant Ranjan, Parimal Das

## Abstract

Calcimycin (A23187) is a polyether antibiotic and divalent cation ionophore, extracted from *Streptomyces chartrecensis*. With wide variety of antimicrobial activities, it also exhibits cytotoxicity of tumor cells. Calcimycin exhibit therapeutic potential against tumor cell growth; however, the molecular mechanism remains to be fully elucidated. Present study explores the mechanism of calcimycin-induced apoptosis cancer cell lines. Calcimycin induces apoptosis accompanied by increased intracellular calcium-level and increased expression of purinergic receptor-P2RX4, a ligand-gated ion channel. The percentage of apoptotic cancer cells in a dose-dependent manner quickly rose as recorded with MTT assays, Phase contrast imaging, wound healing assay, fluorescence imaging by DAPI and AO/EB staining and FACS. Mitochondrial potential was analyzed by TMRM assay as Ca^2+^ signaling is well known to be influenced and synchronized by mitochondria also. Calcimycin treatment tends to increase the intracellular calcium level, mRNA expression of ATP receptor P2RX4, and phosphorylation of p38. Blocking of either intracellular calcium by BAPTA-AM, P2RX4 expression by antagonist 5-BDBD, and phospho-p38 by SB203580, abrogated the apoptotic activity of calcimycin. Taken together, these results show that calcimycin induces apoptosis in P2RX4 dependent ATP mediated intracellular Ca^2+^ and p38 MAPK mediated pathway in both the cancer cell lines.

## 1. Introduction

Continuing explosion in cancer cases makes cancer a second leading cause of death & major public health concern globally. A WHO claims that by 2035 approx 24 million new cancer cases and approx 14.5 million cancer-allied deaths would be expected each year in the world (https://www.cancer.gov/research/areas/global-health). Despite chemotherapy, which is a major cancer treatment strategy, the main impediment in cancer therapy is drug resistance, which can be inherent or acquired (Das & Verma, 2020). As a result, much interest and effort have been focused on researching new strategies and approaches for cancer prevention and treatment with increased effectiveness and fewer side effects. Low-toxicity chemo-preventive agents derived from natural sources, with high potential in tumor inhibition, are promising candidates for cancer treatment. One such agent is a calcium ionophore Calcimycin (A23187), extracted from *Streptomyces chartrecensis* known to have antimicrobial properties (Mawatwal et al., 2017). This ionophore is known to increase cytosolic free calcium (Ca^2+^) levels and thereby increase apoptosis in primary rat cortical cells (Sennvik et al., 2001) and Hela cells (Zhu et al., 2007). It is believed that the majority of calcimycin-induced effects are due to its capability ‘depolarize mitochondrial membrane’ which is supported by the fact that calcimycin encourages ROS generation and Nox5 activation (Benedyk et al., 2007; Przygodzki et al., 2005) and promotes apoptosis in a calcineurin-dependent manner (Kajitani et al., 2007). Also, various autophagy markers like autophagy-related gene (Atg) 7, *Atg 3*, *Beclin-1*, and *LC3-II* are up-regulated in macrophages when exposed to calcimycin (Mawatwal et al., 2017). Although intracellular Ca^2+^ elevation is known to trigger apoptotic cell death (Mu et al., 2008), yet the mechanism of action for calcimycin-induced apoptosis is not fully understood. Kinases such as extracellular signal-regulated kinase (ERK), p38 MAPK, and c-Jun NH (2)-terminal kinases (JNKs) too play a regulatory role in apoptotic processes. According to certain studies, extracellular stress signals that causes p38 MAPK phosphorylation and Ca^2+^ increase encourage apoptosis (Hsu et al., 2007). p38 MAP kinases are important regulators in cell invasion, migration, differentiation, and proliferation too (Jiang et al., 2019). A notable opportunity to check tumor metastasis is provided by the major function that p38 MAPK plays in cell invasion and migration in addition to controlling cell proliferation. p38 MAPK has been suggested to be a key player in leukemia, transformed follicular lymphoma, breast cancer, prostate cancer, lung cancer, liver cancer, and bladder cancer (Koul et al., 2013). Therefore, further study is needed to figure out how the p38 MAPK pathway contributes to oncogenesis and calcimycin-mediated cell death. p38 MAPK activation and enhanced calcium uptake in cells have also been reported to be linked to the ligand-gated ion channel P2RX4 (Trang et al., 2009). A ligand-gated ion channel P2X receptors are a group of ATP-gated ionotropic channels including seven known members P2X1 to P2X7 which regulates quick responses to the transmitter ATP in mammalian cells viz. Leukocytes, vascular smooth muscle central and sensory neurons, endothelium, etc (Fountain, 2013). Among seven P2X receptors, P2X4 (P2RX4) is far and widely expressed in central and peripheral neurons (Suurväli et al., 2017). P2RX4 harbors two transmembrane domains and three ATP binding sites in extracellular loops. Broad admeasurements of P2RX4 in mammalian cells point to the essential regulatory roles of P2RX4 in many biological events. Expression of P2RX4 in numerous cancers such as colorectal, lung, leukemia, bladder cancer, brain tumor, etc. indicates an association of P2RX4 in tumorigenesis, However, P2RX4’s involvement in cancer progression remains to be clarified. P2RX4 is involved in epithelial-to-mesenchymal transition (EMT) and *TGF*β*-1* induced invasiveness in pancreatic cancer (He et al., 2020). These studies allude to the interlink between cancer and P2RX4. Here we comprehensively elaborate the anti-proliferative pathway of calcimycin over cancer cell line via mitochondrial calcium influx mediating caspase-regulated apoptotic cell death. Also, try to link the association of P2RX4 with calcium influx in calcimycin-mediated apoptotic cell death.

## 2. Materials and Methods

### 2.1 Materials

ATP measurement kit was purchased from Promega. Annexin V apoptosis detection kit and western blotting detection reagent ECL were purchased from Thermo Fisher Scientific. Antibiotic cocktail penicillin-streptomycin and MTT (3-(4,5-dimethylthiazol-2-yl)-2,5-diphenyltetrazolium bromide) were purchased from Himedia. DNaseI was purchased from Ambion,Waltham, MA. Calcimycin, p38 MAPK inhibitor SB203580, P2RX4 antagonist 5-BDBD, 5-Fluo-3/AM, TRIzol reagent were purchased from Sigma Aldrich. Calcimycin, SB203580 and 5-BDBD was dissolved in DMSO as stock solution. Final concentration of DMSO was 0.1%.

### 2.2 Cell culture and treatment

Human cervical and breast cancer cell lines SiHa and MCF7 respectively was maintained in Dulbecco’s Modified Eagle’s Medium (DMEM) medium with 10% FBS and 1X penicillinstreptomycin cocktail inside the humidified condition with 5% CO2 at 37°C. For each experiment cells were cultivated for 1 day up to 60-70 % confluence after being seeded from 80-90% confluent T25 flask in complete DMEM media. 60-70 % confluence cells were treated with streptomycin cocktail inside the humidified condition with 5% CO2 at 37°C. For each experiment cells were cultivated for 1 day up to 60-70 % confluence after being seeded from 80-90% confluent T25 flask in complete DMEM media. 60-70 % confluence cells were treated with different concentrations (IC50 and ±10 IC50) of calcimycin for 24 hrs (hours). SiHa was treated with 0.25, 0.35, and 0.45 µM whereas MCF7 was treated with 0.20, 0.30, and 0.40 µM of calcimycin. Additionally for exploring signaling pathways, cells were pretreated with 10 μM BAPTA-AM, 1 μM 5-BDBD, and 10 μM SB203580 for 3 hrs, 1 hr, and 2 hrs respectively followed by incubation with calcimycin at IC50 concentration for 24 hrs.

### 2.3 MTT Assay

Cytotoxicity of the compound was studied by MTT assay according to Verma *et al*., 2018. SiHa and MCF7 cells were seeded in a 96-wells plate at a density of 2×10^4^ with complete DMEM medium inside a 5% CO2 humidified incubator. Next day cells were treated with different concentrations of calcimycin (0 µM to 2 µM). Following 24 hrs (hours) incubation period, media were replaced with DMEM with 0.5 mg/ml MTT and incubated at 37°C. After 4 hrs MTT solution was removed and 100 µl DMSO was added and incubated for 10 minutes in dark to solubilize the blue/purple color formazan. Accumulation of formazan indicates the activity of mitochondria, which indirectly determines the viability of the cell (Riss et al., 2016). The end product of purple color was quantified by measuring absorbance at 570 nm in a microplate reader (Bio-Rad) and the IC50 dose was calculated by dose-response curve. % cell death was calculated by using following formula:

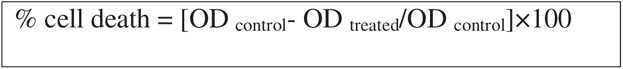

### 2.4 Cellular Morphology assessment

Both cells were seeded in a 6-well plate. After 24 hrs, at 60-70% confluence, cells were treated with calcimycin at IC50 concentration. Images were captured at two-time points (0 and 48 hrs) using a phase-contrast microscope (LEICA) to analyze the morphological effect of the compound on cells.

### 2.5 Wound Healing Assay

Using a 200 μl sterile micropipette tip, the wound was made in monolayer confluence cells. Displaced cells were then removed by rinsing the area with 1X PBS. Cells were treated with IC50 concentrations of calcimycin. Cell migration in the scraped area was assessed by a phase-contrast microscope. At 0, 24, and 48 hrs, pictures were taken. The following formula was used for calculating the wound closure percentage:

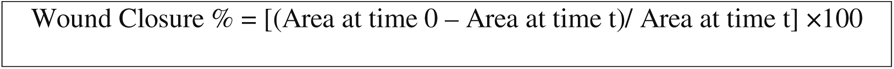

### 2.6 Intracellular Ca^2+^ Measurement

After treatment of cells with calcimycin along with inhibitors, cells were harvested and washed with PBS then stained with 5 μM of Fluo-3/AM for 1 hr in dark at 37°C. Ca^2+^ dependent fluorescence intensity of Fluo-3/AM was measured by flow cytometry (CytoFLEX LX Beckman Coulter).

### 2.7 ATP Measurement

Amount of ATP release in response to compound were measured by Celltiter-Glo 2.0 cell viability assay kit (Promega) as per the manufactures’ protocol. In brief equal no. of cells was grown in white opaque 96 wells plates. Cells were treated with calcimycin as previously described concentration (IC50 and ±10 IC50) for 24 hrs. After the incubation period 100 μl of ready to use celltiter glo reagent were added in each well with media. Contents were mixed for 2 min at orbital shaker to induce the cell lysis following 10 minutes incubation at room temperature luminescent signal was recorded in the illuminometer.

### 2.8 Acridine Orange/Ethidium Bromide (AO/EB) Staining

Morphological evidence of apoptosis was analyzed by AO/EB staining. Calcimycin and corresponding inhibitor’s treated cells were collected by trypsinization followed by PBS washing. After washing cells were stained with 1µg/ml of AO-EB for 5 min at room temperature and images were captured under fluorescence microscope (Evos FLc, Invitrogen).

### 2.9 DAPI Staining

Chromatin condensation was analyzed by DAPI staining; in brief cells were grown in a 12-wells plate on a flame-sterilized cover slip. After the treatment with calcimycin or corresponding inhibitors, cells were washed with PBS and fixed with 4% paraformaldehyde for 15 min at room temperature. After washing twice with PBS cells were permeabilized with 0.5% Triton-X-100 in PBS for 30 min at room temperature. Following permeabilization, cells were washed twice with PBS and stained with 1 µg/ml DAPI for 15 min at room temperature in the dark. After incubation, cells were washed twice with PBS and examined under a fluorescent microscope.

### 2.10 Apoptosis assay by flow cytometry

Apoptosis was evaluated by AnnexinV-APC/PI double staining. After treatment, both cells were harvested by pellet down at 3000 rpm for 3 min and washed twice with PBS. Cells were resuspended in 1X binding buffer and stained with 5 μl Annexin V-APC and 1 μl PI and incubated for 15 min at 37 °C in the dark. Measurements were performed by a flow cytometry (CytoFLEX LX Beckman Coulter). 10,000 events were counted for each sample.

### 2.11 Cell Cycle Analysis

Cells were stained with PI to study the different phases of the cell cycle as a result of calcimycin treatment. After treatment with calcimycin at IC50 concentration, cells were harvested at 3000 rpm for 3 min and then washed with PBS. Cells were fixed in cold 70% ethanol for 45 min at −20 °C. After fixation, the cells were rinsed in PBS, and the pellet was then resuspended in 200 μl of PBS containing 50 μg/μl of PI and 50 μg/μl of RNase, which was then incubated for 30 min at room temperature in the dark. Stained cells were analyzed by flow cytometry (CytoFLEX LX Beckman Coulter).

### 2.12 TMRM Assay

The effect of calcimycin on mitochondrial membrane potential was measured by a TMRM fluorescent dye. Cells were harvested after incubation with calcimycin for 24 hrs followed by washing with PBS and further stained with 200 nM TMRM in PBS at 37°C for 15 min in the dark. Flow cytometry was performed to acquire orange fluorescence intensity.

### 2.13 Gene Expression Analysis by qRT-PCR

Drug effect on the expression of genes related to cell death was analyzed by quantitative real-time PCR (qRT-PCR) according to Verma *et al*., 2018. In brief total cellular RNA was extracted from both treated and non-treated (control) cells by using TRIzol reagent followed by DNaseI treatment to remove DNA contamination. The concentration and purity of RNA were checked by NanoDrop (Thermo, Waltham, MA). After that cDNA was synthesized from 1µg of purified RNA by using a cDNA synthesis kit (Applied Biosystems, Waltham, MA) as per the manufacturer’s instructions. The expression levels of selected genes were determined using qRT-PCR (QuantStudio 6, ThemoFisher Scientific) using the SYBR Green Real-time PCR master mix (Thermo Scientific). The sequence of the primers used for the gene expression analysis was given in an appendix.

### 2.14 Western Blotting

Cells were trypsinized after calcimycin treatment (together with inhibitors), harvested, and washed in cold 1XPBS before being pelleted at 4000 rpm for five minutes at 4°C. Following a 30-minute incubation on ice in RIPA buffer, the cell pellet was lysed. Protein lysate was collected by centrifugation at 13000 rpm for 30 min at 4°C. Protein quantification was done by Bradford reagent. An equal amount (40 µg) of protein was loaded and resolved on the 10% SDS-PAGE. Specific proteins were detected by their corresponding antibodies, and β actin was used for loading control.

### 2.15 Docking

As the structures of human P2RX1, P2RX2, P2RX4, P2RX5, P2RX6, and P2RX7 were unavailable in the Protein Data Bank (Berman et al., 2007), approaches of homology modeling prediction using the Swiss model (Schwede et al., 2003) were used to derive the 3D structure of the P2RX receptor family. The template used for the prediction of the P2RX receptor family structure was 5SVK. However, the crystal structure of P2RX3 (human) was available in PDB (5SVK) and could be retrieved and used for further processing of the docking. The selection of template structures for P2RX1, P2RX2, P2RX4, P2RX5, P2RX6, and P2RX7 (NCBI Reference Sequence: NP_002549.1, AAF74204.1, NP_001243725.1, BAD92860.1, NP_005437.2, and CAA73360.1) was determined based on parameters of maximum score and query coverage. Modrefiner (Xu & Zhang, 2011) was used to refine the derived structures. The quality of the derived structures was further validated by the PDBsum server (Laskowski et al., 2018).The 3D structures thus generated were visualized by Discovery Studio 4.0 (Studio, 2008). The CASTp 3.0 server (Tian et al., 2018) predicted the active binding sites for the chosen 3D structures and identified the active site residues. Discovery Studio 4.0 was used for further visualization of active pockets of the structures. The PubChem Compound database (Kim et al., 2019) was used to retrieve the 3D structure of the ligand calcimycin (CID 40486), which was then converted from SDF to PDB format and optimized using Discovery Studio 4.0. By using PatchDock Server (Schneidman-Duhovny et al., 2005) (parameter RMSD value 1.0 and complex type protein-small ligand), calcimycin was docked with several chosen targets. Discovery Studio 4.0 was then used to visualize the docked complexes. Using the Prodigy server (Xue et al., 2016), binding energy (BE) was calculated. Additionally, the docked complex was used in the Prodigy server to calculate binding energy. More negative energies suggest a stronger bond.

### 2.16 Statistical Analysis

Each experiment was performed three times, and the data were provided as the mean standard deviation (Standard Deviation). The statistical analysis was performed using the paired student t-test. When the P-value was ≤ 0.05, the differences were considered statistically significant.

## 3. Results and Discussion

### 3.1 Antiproliferative effect of calcimycin

The MTT test was used to examine and analyze the antiproliferative activity of calcimycin against cervical and breast cancer cell lines. Cell viability analysis was carried out with varying concentrations of calcimycin (0 to 2 μM) and revealed dose-dependent growth inhibition activity, as approx 90% cell death was achieved at 2 μM concentration, indicating that the calcimycin inhibits cell survival and proliferation (Fig-1). IC50 value after 24 hrs treatment of calcimycin was found to be 0.35µM & 0.30µM for SiHa and MCF7 cell lines respectively. Changes in cellular morphology in response to calcimycin at IC50 dose were studied by phase contrast microscopy. Both SiHa and MCF7 cells were found to become vacuolated, detached from the surface, and floating in the medium after the 24 hrs treatment of calcimycin as compared to control cells (untreated) shown in Fig-2. The phase contrast images demonstrated that calcimycin caused cell shrinkage and detachment from the surface of the monolayer while the reduction in the number of viable cells, supporting the findings of the MTT cell viability assay.

**Figure 1:**
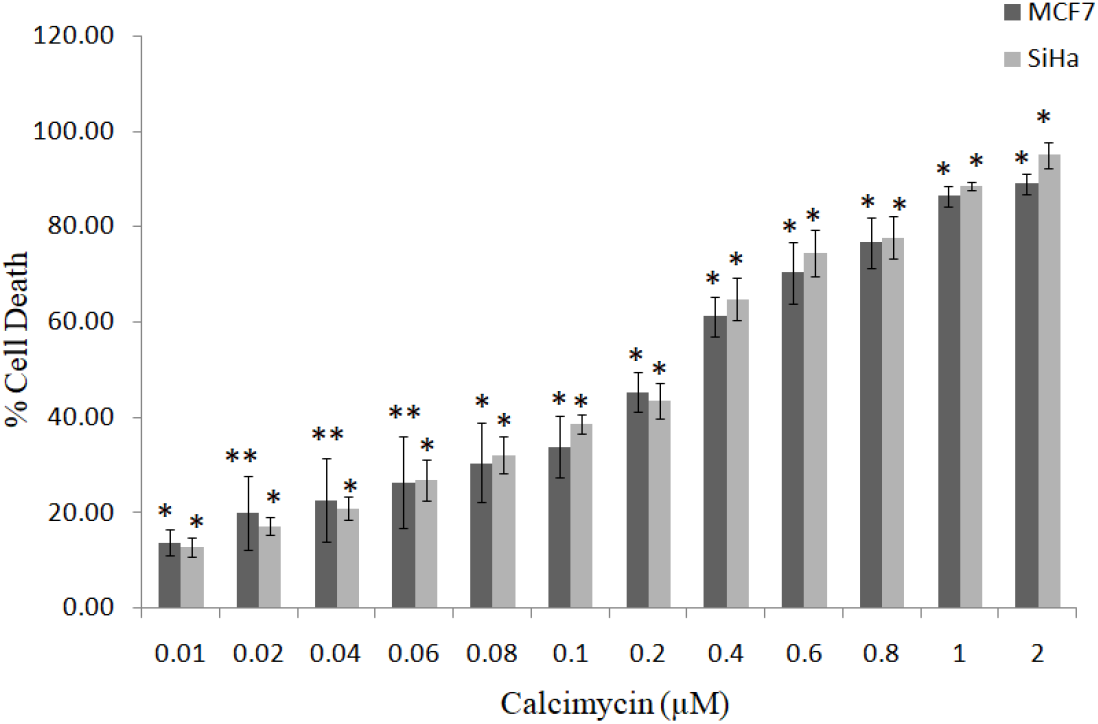
Antiproliferative effect of calcimycin in SiHa and MCF7 cell lines. Calcimycin induces cell death in dose (0.01-2µM) dependent manner. Light grey and dark grey bar showing % cell death at 24 hrs in SiHa and MCF7 respectively. The error bar shows the mean and SD. The paired t-test was used to measure statistical significance. *P ≤ 0.05, **P ≤ 0.01 compared with control vs. other groups

**Figure 2:**
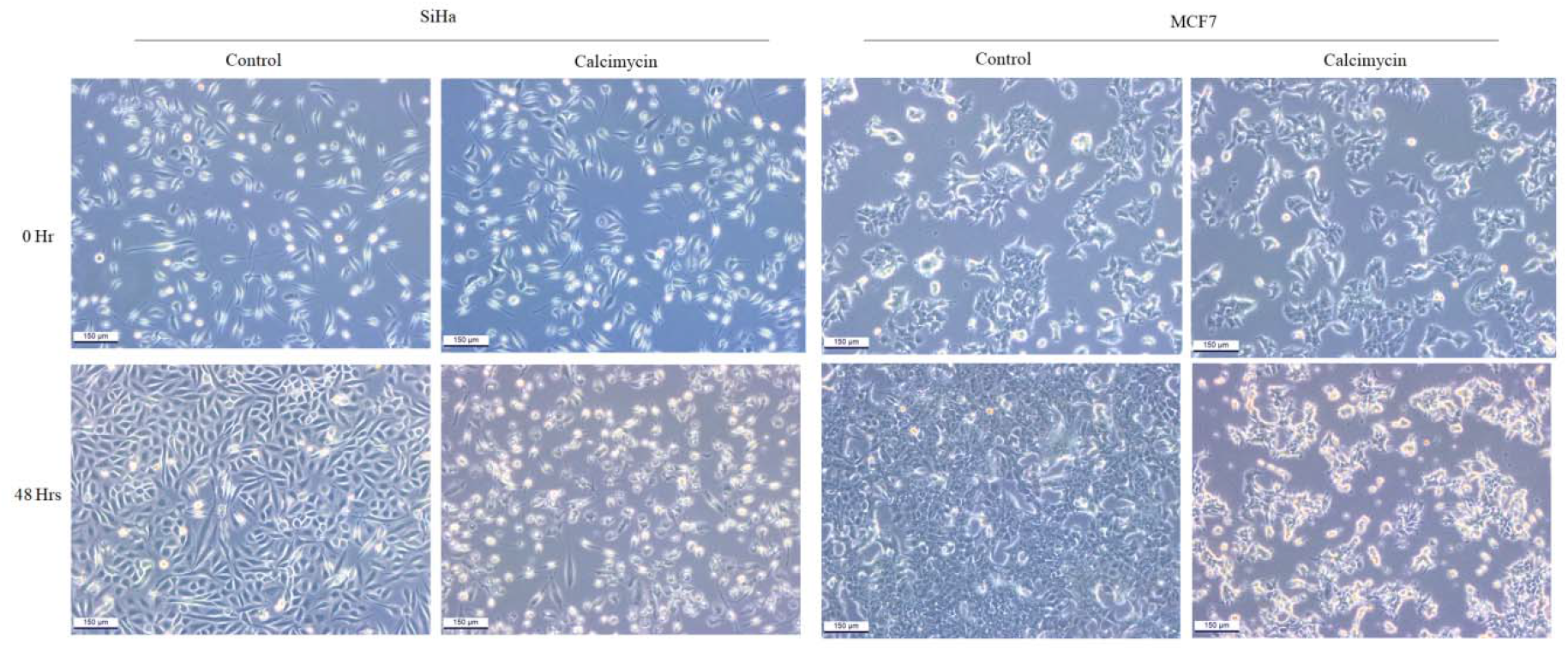
Phase contrast microscopy shows cell death induced by calcimycin. After 48 hrs of incubation with calcimycin of 0.30 µM in MCF7 & 0.35 µM in SiHa, cells became rounded, vacuolated, and floating in the medium where as in control (no treatment) no cell death was observed and cells achieved their confluency at 48 hrs. Scale bar is 150 μm

### 3.2 Calcimycin inhibits cell migration

A wound healing assay was performed to observe cell migrations towards the wounded area after incubation with or without calcimycin at the IC50 dose at three different time points viz., 0, 24, and 48 hrs (Fig-3). The calculation of the wound closure percentage was done by examining the change in scratch spacing over time. As shown in Fig-3 untreated SiHa had a wound size of 297.18 nm at 0 hr, which decreased to 170.85 nm at 24 hrs and 56.16 nm at 48 hrs, whereas treated SiHa had a wound size of 388.4 nm at 0 hr, which decreased to 300.69 nm at 24 hrs and 265.61 nm at 48 hrs. Similarly, untreated MCF7 wound size was reduced from 403.69 nm at 0 hr to 222.32 nm at 24 hrs and 92.43 nm at 48 hrs, whereas treated MCF7 wound size was reduced from 437.60 nm at 0 hr to 373.23 nm and 303.03 nm at 24 hrs and 48 hrs, respectively. Wound closure percentage in untreated SiHa was found 73.9 and 429.16% at 24 and 48 hrs respectively while in the case of treated SiHa, it was 29.16 and 46.22 % at 24 and 48 hrs respectively. Untreated MCF7 wound closure percentage is 81.5% at 24 hrs and 336.7% at 48 hrs, whereas treated MCF7 wound closure percentage is 14.7% at 24 hrs and 44.4% at 48 hrs. In both SiHa and MCF7 cells, calcimycin treatment highly reduced cell migration activity compared to control/untreated cells.

**Figure 3:**
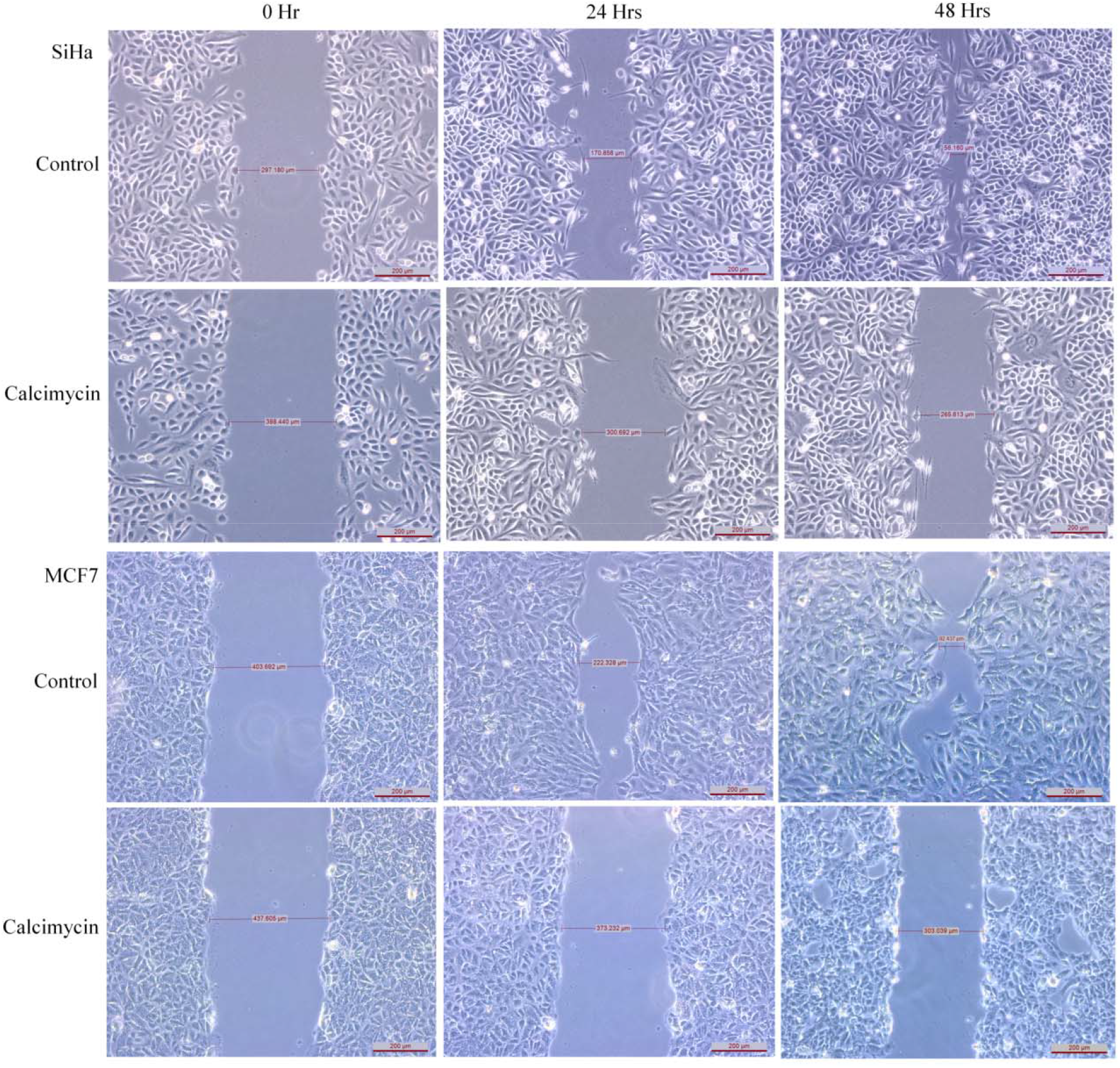
Wound Healing assay shows calcimycin inhibits cellular migration in SiHa and MCF7 cell lines. After 24 and 48 hrs incubation with calcimycin of 0.30 µM in MCF7 & 0.35 µM in SiHa, scratch gap within the cells persist while in control gaps get refilled by cell migration. Scale bar is 200 μm

### 3.3 Effect of calcimycin on cell apoptosis

To record morphological alteration, Chromatin condensation & nuclear fragmentation to demonstrate apoptotic cell death both SiHa and MCF7 cells were stained with DAPI after incubation with calcimycin. In calcimycin-treated cells, DAPI staining revealed increased chromatin condensation and nuclear fragmentation in a dose-dependent manner (Fig-4). To further validate apoptotic cell death, cells were stained with AO/EB staining. The number of EB-stained nuclei increased in a dose-dependent manner confirming the presence or increment of early and late apoptosis cells in treated conditions (Fig 5). Further, annexin V-APC/PI assay was employed to evaluate apoptosis. Flow cytometry analysis of both cell lines shows enhancement of apoptotic cell populations with increasing concentration of calcimycin as compared to control cells. In SiHa after 24 hrs treatment of the test compound late apoptotic cells were 6.15%, 8.21%, and 10.98% while early apoptotic cells were 14.87%, 15.99%, and 5.99 % at 0.25, 0.35, and 0.45µM concentration respectively as compares to control cells having 3.53% early and 4.29% late apoptotic cells. Similarly, in MCF7 late apoptotic cells were 4.64%, 7.82%, and 13.64% while early apoptotic cells were 4.77%, 3.80%, and 3.93% at 0.20, 0.30, and 0.40µM concentration respectively compare with control 2.36% early and 3.41% late apoptotic cells as shown in Fig-6. In this study, calcimycin was found to be cytotoxic to breast and cervical cancer through apoptosis in a dose-dependent manner.

**Figure 4:**
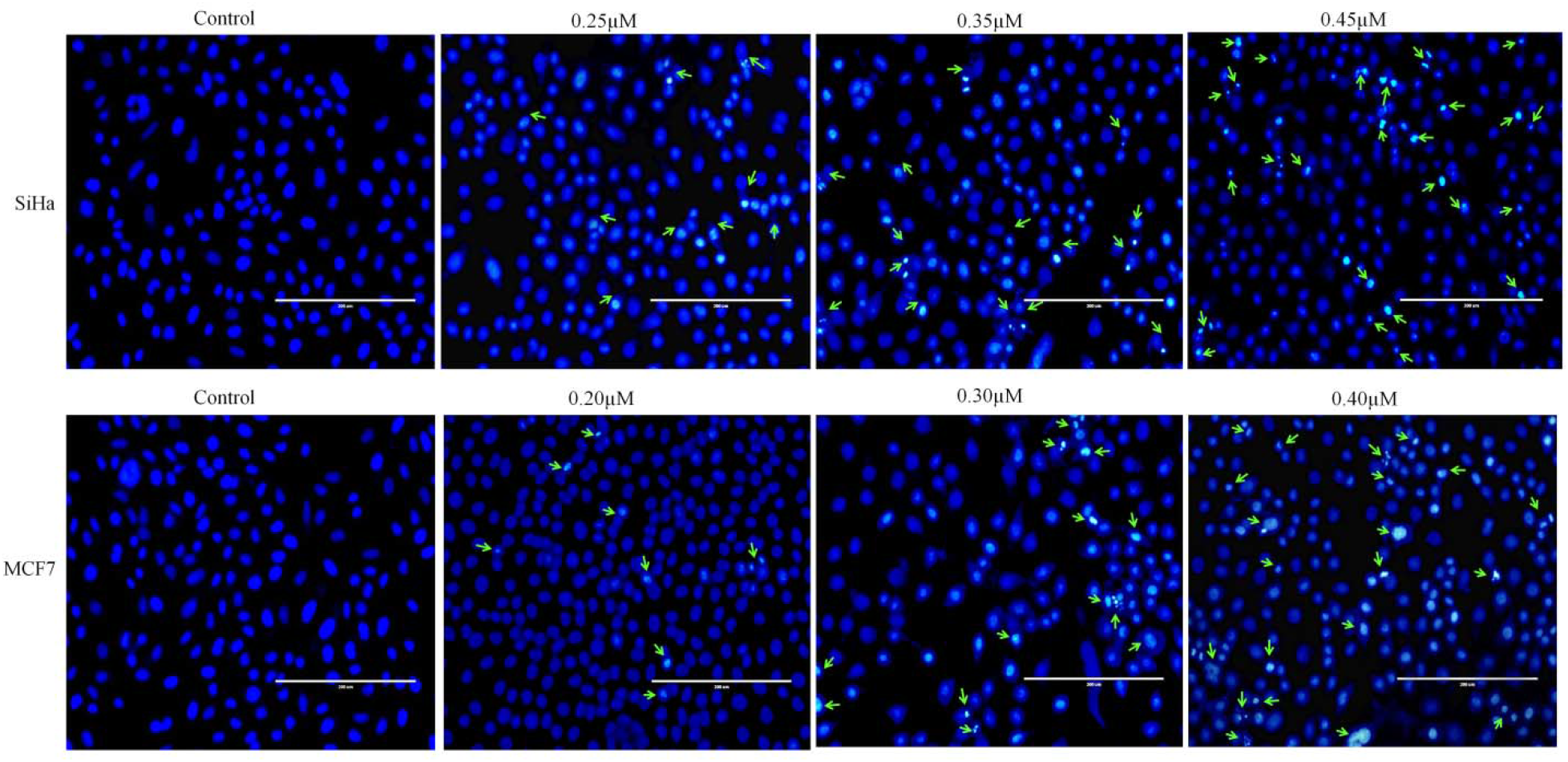
Nuclear staining by DAPI of SiHa and MCF7. Arrows indicate nuclear fragmentation, cell shrinkage and margination of the nucleus which increases with increasing concentration of calcimycin in comparison to control cells. Scale bar is 200 μm

**Figure 5:**
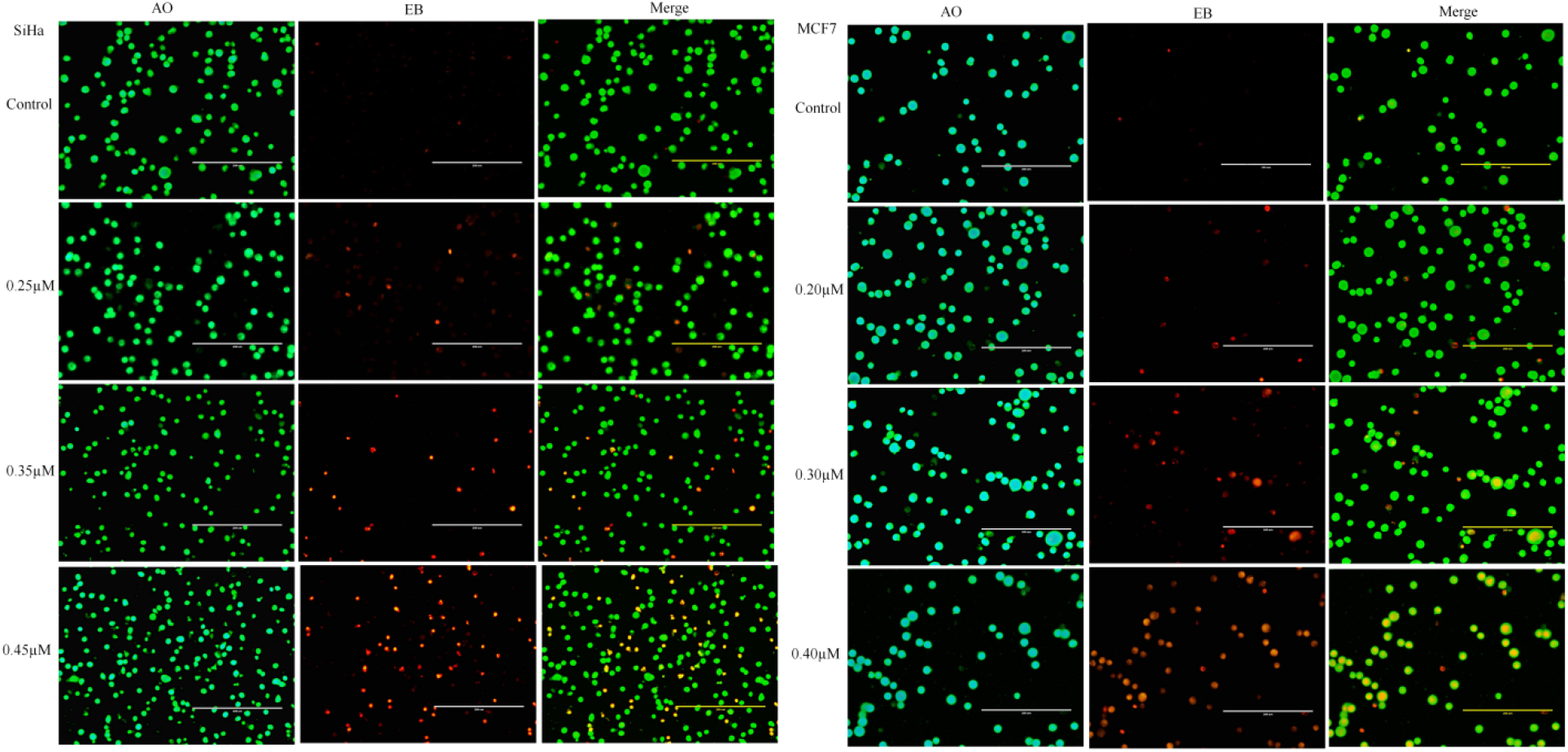
Detection of apoptosis by morphological changes using acridine orange and ethidium bromide (AO-EB) staining in SiHa and MCF7. Cells stained green are living, early & late apoptosis is represented by yellow/orange colored. No. of early & late apoptotic cells was increases in treated cells in dose dependent manner. Scale bar is 200 μm

**Figure 6:**
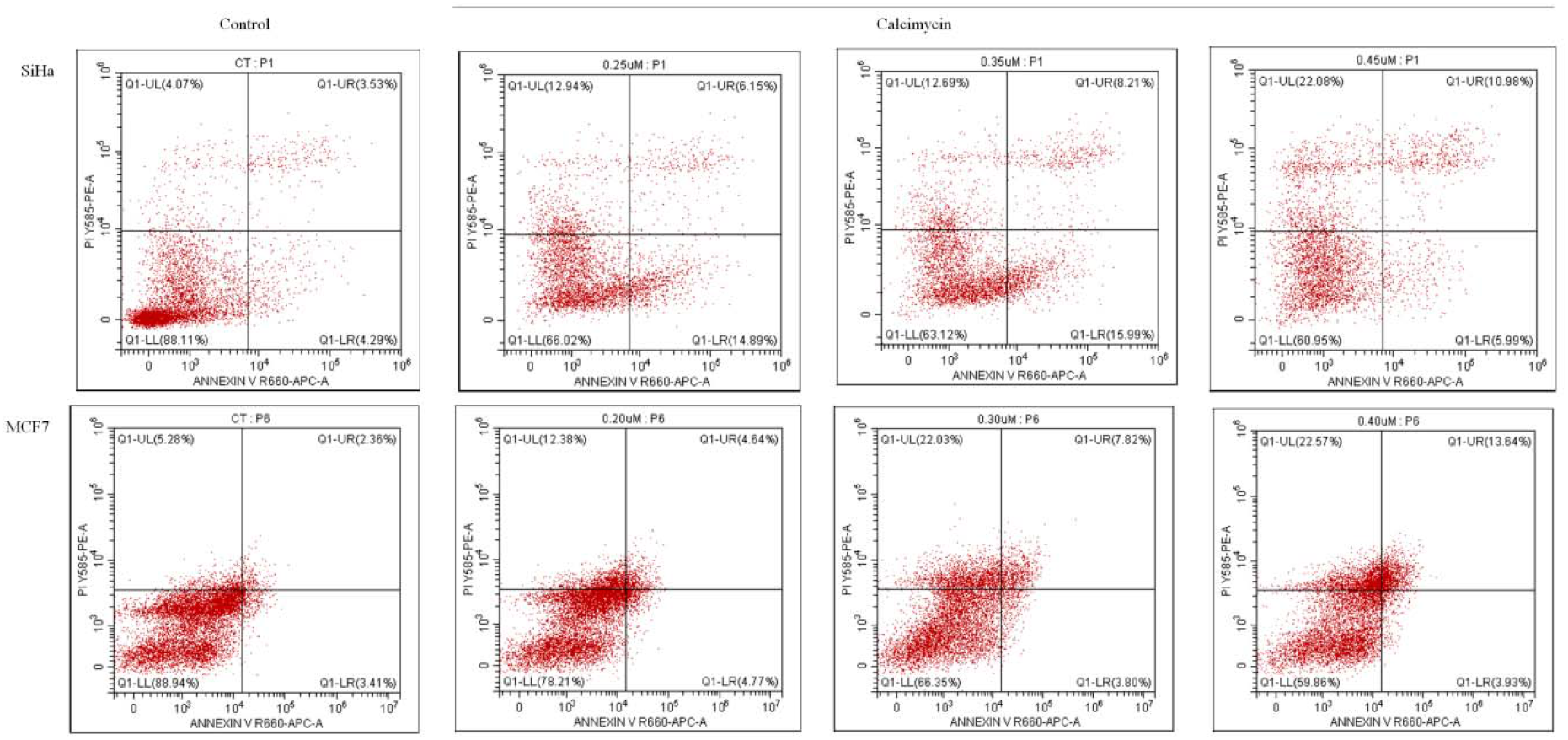
Apoptosis evaluation in SiHa and MCF7 cell lines by flow cytometry by Annexin V-APC and PI staining. No. of late apoptotic cells were increased with increasing concentration of calcimycin as compared to control cells

### 3.4 Calcimycin induces cell cycle arrest

Next, we tried to investigate cell cycle arrest in response to calcimycin. Different stages of the cell cycle viz. Sub G0, G1/G1, S, and G2/M were plotted and analyzed by DNA histogram. Both SiHa and MCF7 show a significant increase in cell number at the G2/M phase from control 19.20% and 26.01% to treated 38.29% and 68.24% respectively. An increase in peak for the G2/M phase signifies calcimycin causes cell cycle arrest at the G2/M phase with a proportional decrease in cell population in sub-G0, G0/G1, and S phase compared with control (Fig 7).

**Figure 7:**
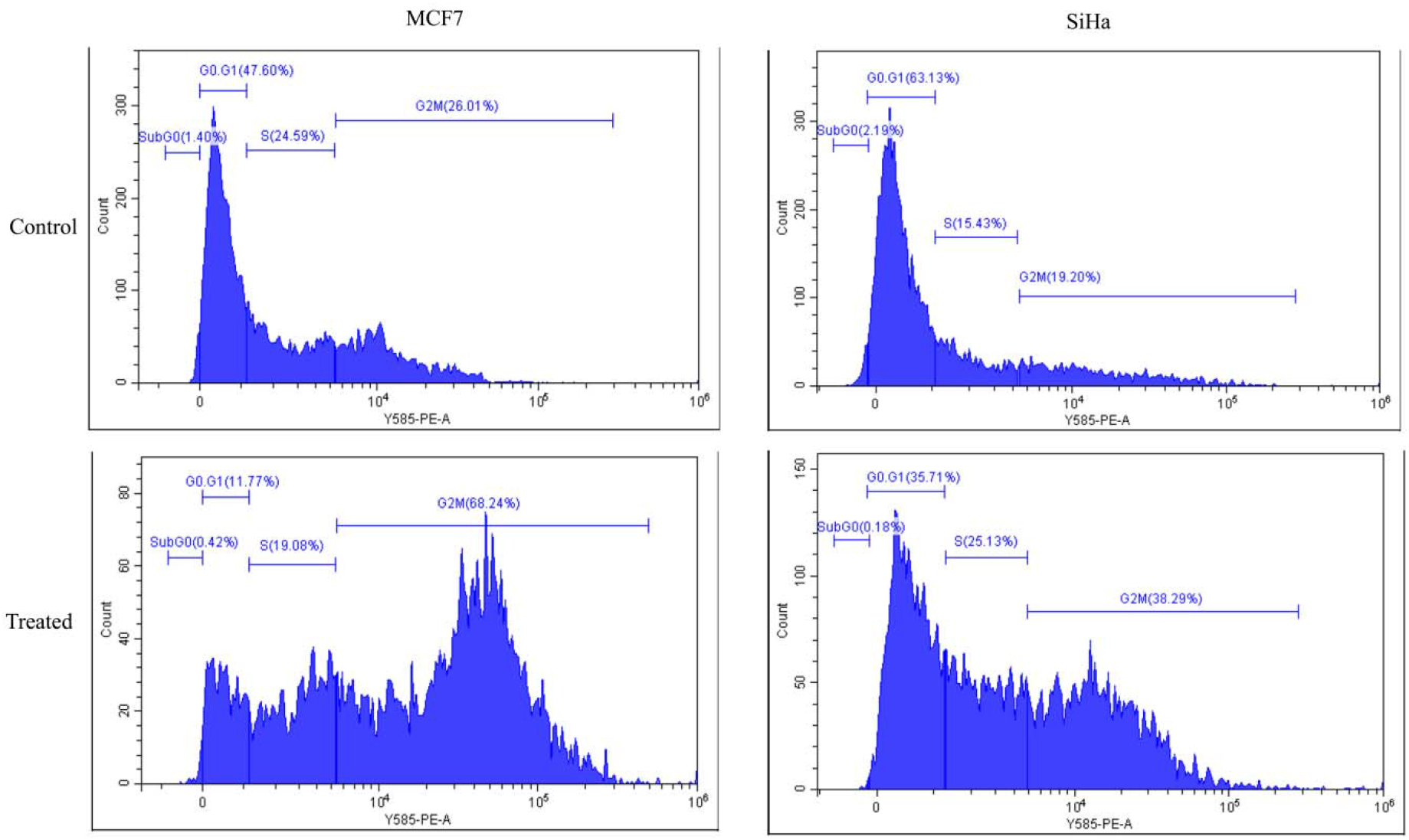
cell cycle analysis of SiHa and MCF7 following calcimycin treatment. Calcimycin treatment of 0.30 µM in MCF7 & 0.35 µM in SiHa induced G2M phase arrest of the cell cycle as compared to control

### 3.5 Role of Intracellular Ca^2+^ level in calcimycin induced apoptosis

The effect of calcimycin on intracellular Ca^2+^ levels was done by calcium measurement using Fluo-3/AM by flow cytometry. As shown in Fig-8-A peaks corresponding to the intracellular Ca^2+^ get shifted toward the right which indicates the enhancement of intracellular Ca^2+^ with increasing concentration of calcimycin in both cell lines. To further verify the intracellular Ca^2+^ elevation in calcimycin induces apoptotic cell death; cells were pretreated with intracellular Ca^2+^ chelator BAPTA-AM for 3 hrs before 24 hrs incubation of test compound to prevent the Ca^2+^ level enhancement followed by DAPI and AO/EB staining along with apoptosis measurement with annexin V-APC/PI assay. As expected intracellular Ca^2+^ level decreases with the pretreatment of BAPTA-AM (Fig-8-B). In both SiHa and MCF7 nuclear fragmentation and chromatin condensation were found to restore in BAPTA-AM pretreated/ Ca^2+^ chelated cells compared to only calcimycin-treated cells. Annexin V-APC/PI staining also indicated a fall in no. of late apoptotic cells in Ca^2+^ chelated cells i.e 4.83% & 1.64% from calcimycin treated cells i.e. 8.15% & 11.91% in both cell lines SiHa & MCF7 respectively as shown in Fig-9. Hence calcimycin induces the intracellular Ca^2+^ increase and apoptotic events which get attenuated by chelating intracellular Ca^2+^ in both breast and cervical cancer cell lines.

**Figure 8:**
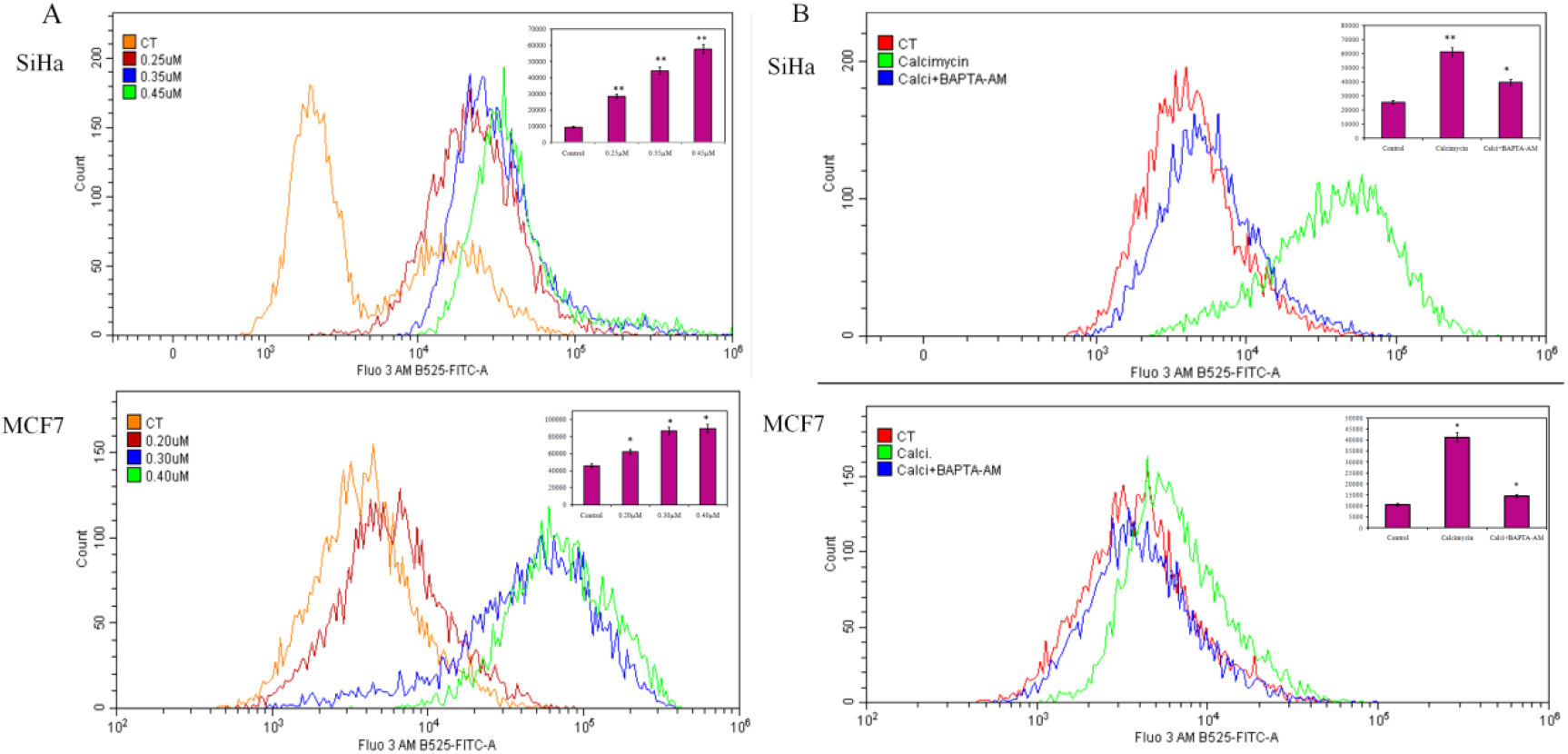
(A) calcimycin treatment induces increament of intracellular Ca^2+^ level in SiHa and MCF7 cell line in dose dependent manner. Peak shifted toward right side. Control (orange), <10 IC50 (red), IC50 (blue), >10 IC50 (green). (B) Pretreatment with intracellular Ca^2+^ chelator BAPTA-AM reduces intracellular Ca^2+^ level. Control (Red), only calcimycin (green), calcimycin+BAPTA-AM (blue). In inset graph showing the significant change in Ca^2+^ level. The error bar shows the mean and SD. The paired t-test was used to measure statistical significance. *P ≤ 0.05, **P ≤ 0.01 compared with control vs. other groups

**Figure 9:**
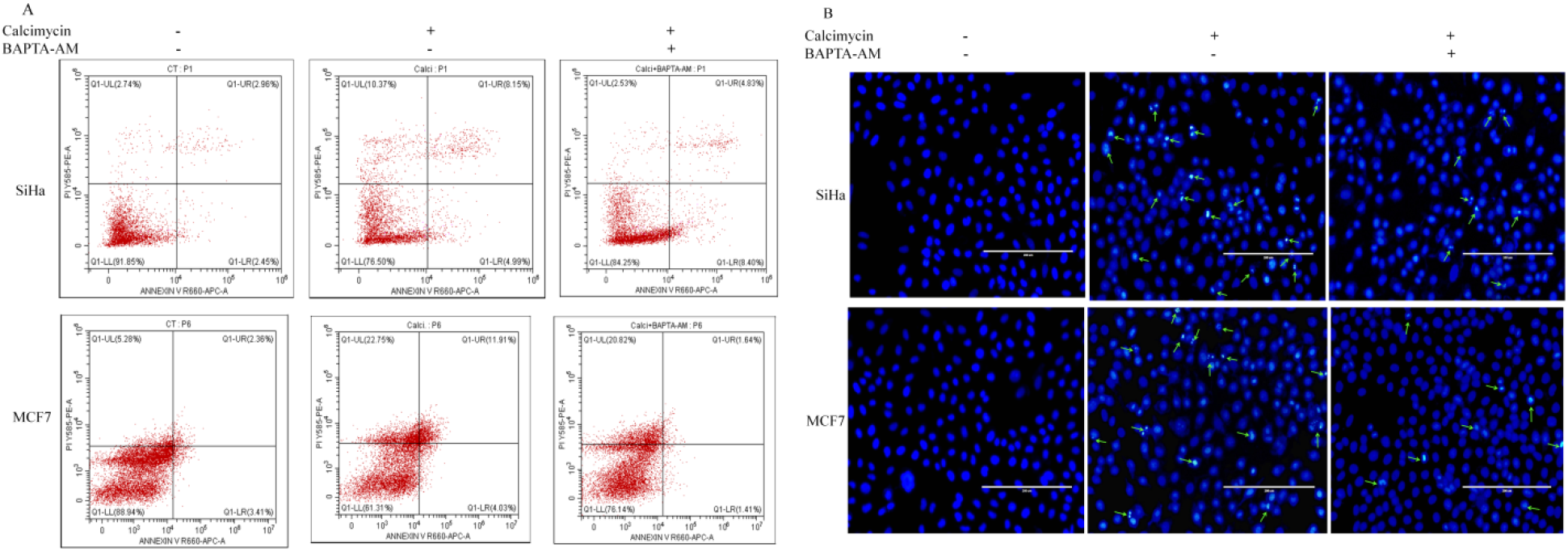
Effect of intracellular Ca^2+^ level in calcimycin induced apoptosis. (A) Effect of calcium chelator BAPTA-AM on apoptosis by flow cytometry. In the effect of BAPTA-AM late/early apoptotic cell no. get reduces in comparison to only calcimycin treated cells. (B) Effect of calcium chelator BAPTA-AM on apoptosis by nuclear stain DAPI staining. Nuclear fragmentation and cell shrinkage also tends to reduce in BAPTA-AM+calcimycin double treated cells compared to single treated (calcimycin only) cells. Scale bar is 200 μm

### 3.6 Effect of calcimycin on Mitochondrial membrane potential (**Δψ**m)

Measurement of Δψm was done by fluorescent dye tetramethylrhodamine methyl esters (TMRM) which gives orange fluorescence. As shown in Fig-10, the TMRM intensity peak has shifted toward the left with an increasing concentration of calcimycin indicating significant reduction in fluorescence after calcimycin treatment relative to control. These results signify the reduction in mitochondrial membrane potential in both cell lines by calcimycin treatment suggesting possibility of mitochondrial Ca^2+^ influx in response to calcimycin treatment.

**Figure 10:**
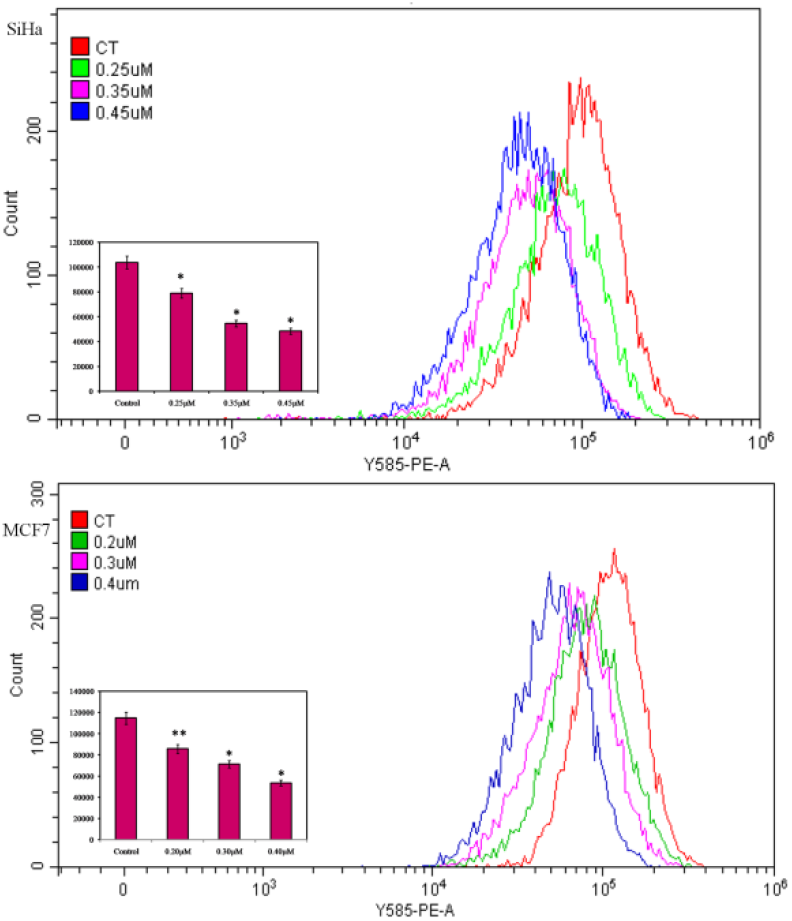
Mitochondrial membrane potential (Δψm) assessment by flow cytometry. TMRM fluorescence get reduced (peak shifted towards left) with increasing concentration of calcimycin in both SiHa and MCF7 cell lines. Control (red), <10 IC50 (green), IC50 (pink), <10 IC50 (blue). In inset graph showing the significant reduction in TMRM fluorescence intensity. The error bar shows the mean and SD. The paired t-test was used to measure statistical significance. *P ≤ 0.05, **P ≤ 0.01 compared with control vs. other groups

### 3.7 Involvement of P2RX4 dependent ATP mediated pathway in calcimycin induced Apoptosis

At first, transcript level expression of P2RX4 in response to calcimycin was evaluated by qRT-PCR. In SiHa we observed 1.52, 2.90, and 4.32 fold increases in P2RX4 mRNA expression with 0.25, 0.35 & 0.45 µM concentrations of calcimycin respectively, similarly noticed 1.29, 2.65, & 4.02 fold increases in MCF7 with 0.20, 0.30 & 0.40 µM calcimycin treatments respectively (Fig-11-B). Next ATP measurements with or without calcimycin treatment were done. Fig-11-C shows enhanced normalized extracellular ATP levels by 1.19, 1.32 & 1.38 fold in SiHa, similarly 1.29, 1.38 & 1.64 fold in MCF7 cells with increasing concentration of calcimycin. Further to elucidate the involvement of P2RX4 in calcimycin-mediated intracellular Ca^2+^ elevation and apoptosis we checked the intracellular Ca^2+^ level with respect to P2RX4 antagonist 5-BDBD. As shown in Fig-11-A 1 hr prior treatment of 1µM of 5-BDBD followed by 24 hrs treatment of calcimycin results in a decrease in intracellular Ca^2+^ in both SiHa and MCF7 cells, suggesting calcimycin raising intracellular Ca^2+^ through ATP receptor P2RX4. We next checked the apoptotic events by DAPI and AO/EB staining along with apoptosis measurement with annexin V-APC/PI assay in response to P2RX4 antagonist 5-BDBD. As expected, DAPI staining shows nuclear fragmentation and chromatin condensation in both SiHa and MCF7 decreases in 5-BDBD pretreated cells compared to only calcimycin-treated cells in Fig-12-B. Likewise, AO/EB staining also showed reduced no. of red-colored early-late apoptotic cells population in 5-BDBD pretreated cells in comparison to only calcimycin-treated cells (Fig-12-A). Further Annexin V-APC/PI staining shows a reduction in no. of late apoptotic cells in 5-BDBD pretreated cells in comparison to only calcimycin treated cells. In SiHa late apoptotic cells were reduced from 8.15% to 5.38% while in MCF7 from 7.93% to 1.27% from without inhibitor to with inhibitor respectively as shown in Fig-12-C. Further, we carried out a molecular docking study to find out how calcimycin affects intracellular calcium levels via P2RX4. After refinement of structure (via modrefiner), the targets were evaluated by pdbsum, and found that more than 90 percent of the residues fell under the most favorable regions in the Ramachandran plot, which showed good structure for docking analysis. Calcimycin docked with the structure of all the 7 members of the P2RX receptor family and found the highest binding energy (−8.5 Kcal/mol) against P2RX4 as compared to other receptors (Fig 13-A), supporting our wet lab data. Calcimycin has a high binding affinity for P2RX4 at its active pocket. The molecular interaction of calcimycin with P2RX4 was depicted in Fig-13 and the hydrogen bonding of calcimycin with the active site residue of P2RX4 is shown in Table 1. These findings suggest that calcimycin binding to P2RX4 contributes to the elevation of intracellular calcium levels.

**Figure 11:**
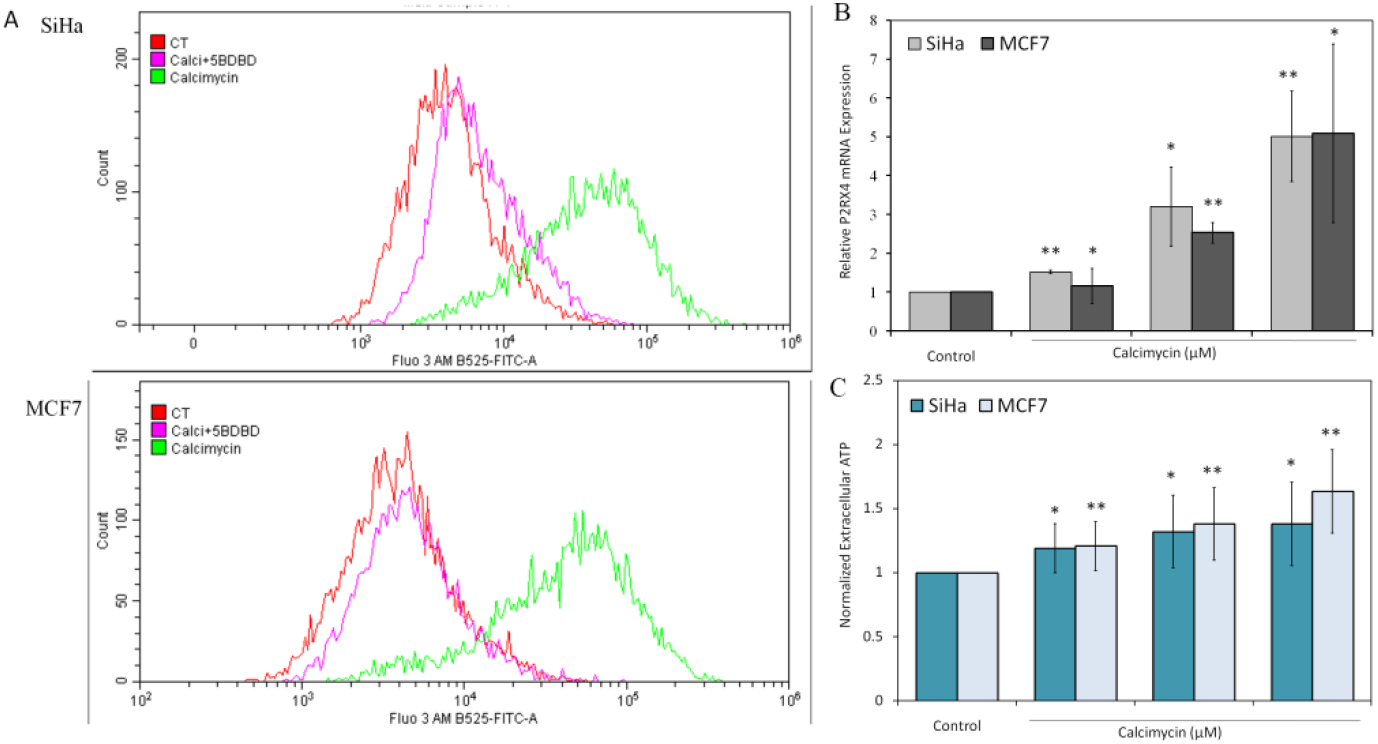
(A) Effect of 5-BDBD upon intracellular Ca^2+^ level in SiHa and MCF7 cell line. In the double positive cells i.e 5-BDBD+calcimycin (pink), intracellular Ca^2+^ gets reduced as compared to calcimycin treatment only (green). Control (red). (B) Effect of calcimycin over P2RX4 mRNA expression by qrtPCR. With increasing concentration of calcimycin, relative mRNA expression of P2RX4 gene get increases also in both cell lines. (C) Effect of calcimycin over extracellular ATP level. Bar of normalized extracellular ATP get increases with increasing concentration of calcimycin in SiHa (light blue) and MCF7 cells (white).The error bar shows the mean and SD. The paired t-test was used to measure statistical significance. *P ≤ 0.05, **P ≤ 0.01 compared with control vs. other groups

**Figure 12:**
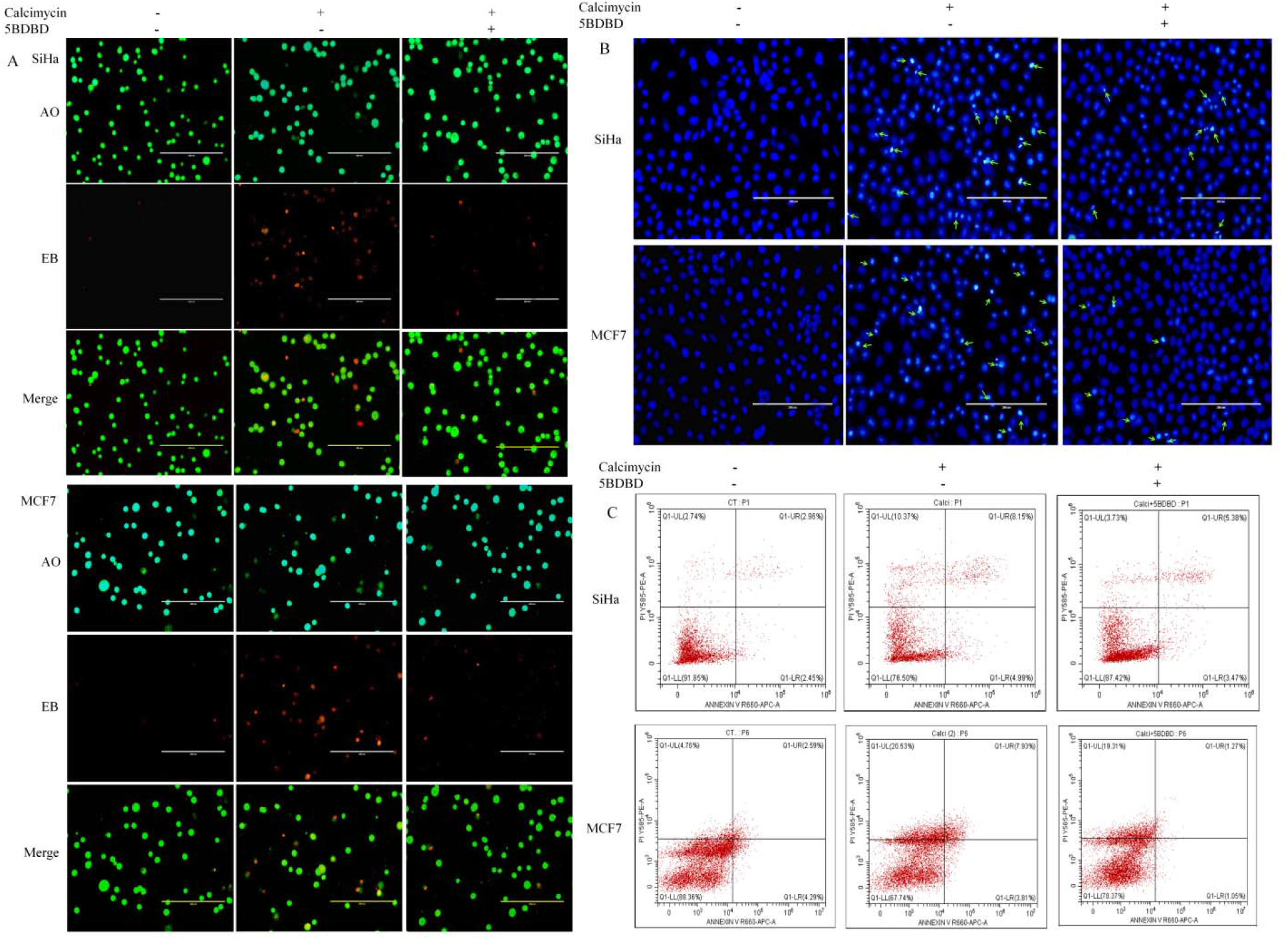
Effect of P2RX4 antagonist 5-BDBD in calcimycin induced apoptosis. (A) Effect of 5-BDBD on apoptosis by AO/EB staining. Early/late apoptotic cell no. get reduced in double positive cells i.e 5-BDBD+calcimycin in comparison to calcimycin treatment only in both SiHa and MCF7 cell lines. (B) Effect of 5-BDBD on calcimycin mediated apoptosis by nuclear stain DAPI staining. In the effect of 5-BDBD, nuclear condensation, chromatin fragmentation in SiHa and MCF7 get diminished as compared to calcimycin treatment only. Scale bar is 200 μm. (C) Flow cytometry assay shows late apoptotic cell no. get decreases in 5-BDBD+calcimycin treated cells in comparison to only calcimycin treated cells

**Figure 13.**
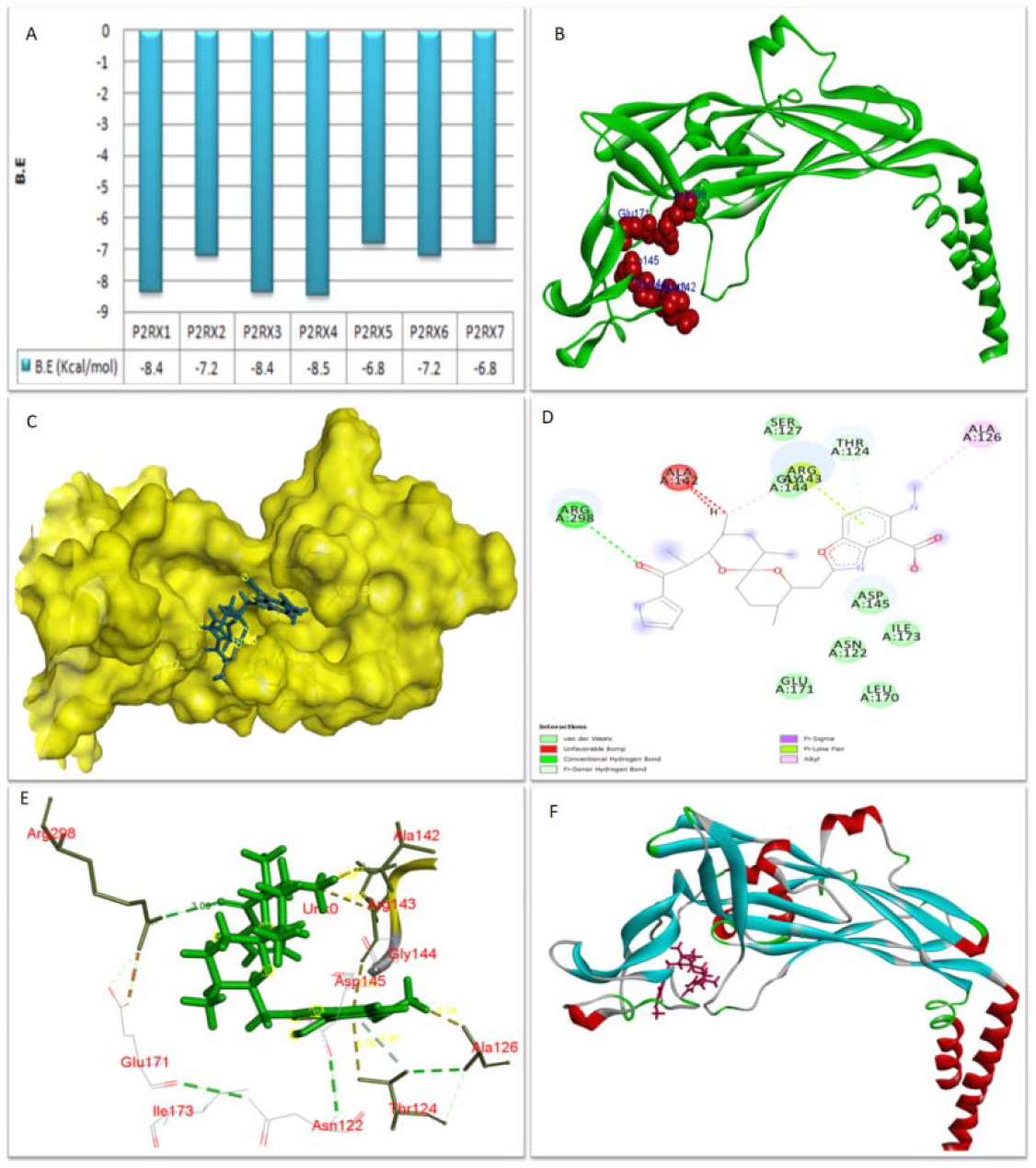
Molecular docking of calcimycin to the family of P2RX receptors. **A.** The results of calcimycin’s molecular docking with the P2RX receptor family were represented graphically, demonstrating the differences in BE (Binding Energy) with regard to calcimycin’s molecular docking with the P2RX receptor family. **B.** Red bubbles indicate the binding pockets in the P2RX4 receptor (green in color). **C.** Calcimycin binds in the cavity of the binding pockets of the P2RX4 receptor. **D and E** represents a 2D and 3D interaction plot of docking results. Hydrophobic interaction was displayed in 2D plots using various colours. **F.** 3D interactions between proteins and ligands

**Table 1:** The representations of active site analysis

| Active sites of P2RX4 | Residues interacts with<br>Calcimycin | Active site residues<br>interacted with calcimycin |
| --- | --- | --- |
| 114GLN, 122ASN,<br>142ALA,143ARG,144GLY,<br>145ASP, 166SER,<br>167TRP,170LEU, 171GLU,<br>298ARG, 312ARG, 314LEU | Asn122, 124Thr, 126ALA<br>,142ALA,143ARG,144GLY,<br>145ASP, 171Glu, 173Ile,<br>298Arg | 142ALA,143ARG,144GLY,<br>145ASP, 171GLU, 298ARG |

### 3.8 Role of p38 MAPK in calcimycin mediated Apoptosis

Caspase 3 is a critical mediator of apoptosis. To check the role of calcimycin induced apoptosis at molecular level we analyzed the expression of cleaved caspase 3 in the presence or absence of calcimycin and inhibitors. As expected western blot result showing the activation of cleaved caspase 3 in calcimycin treated cells which get restored in the presence of SB203580, 5-BDBD and BAPTA-AM (Fig-14). Involvement of p38 and phospho p38 (pp38) in calcimycin mediated apoptosis were also done by western blotting. As shown in Fig-14, expression level of phospho p38 was increased in the presence of calcimycin which got reduced in the effect of prior treatment of BAPTA-AM, 5-BDBD and SB203580, however no change was observed in p38 expression level. In continuation with our study next we examined whether the apoptotic stimuli caused by calcimycin treatment is p38 MAPK mediated or not. As shown in Fig-15-B, DAPI staining showed reduced no. of cells showing chromatin condensation and nuclear degradation in prior treatment with pp38 inhibitor as compared to only calcimycin treatment. AO/EB staining assay also shows attenuation in cell death as early-late apoptotic cells get decreases in the presence of pp38 inhibitor as compared to calcimycin treated cells (Fig-15-A). These results were consistent with annexin V-APC/PI assay where late apoptotic cell population gets reduced from 9.72% to 4.96% in SiHa and 11.51% to 4.78% in MCF7 from calcimycin treatment to pp38 inhibitor treatment respectively (Fig-15-C), which supports the hypothesis of participation of p38 MAPK pathway in apoptosis induction by calcimycin.

**Figure 14:**
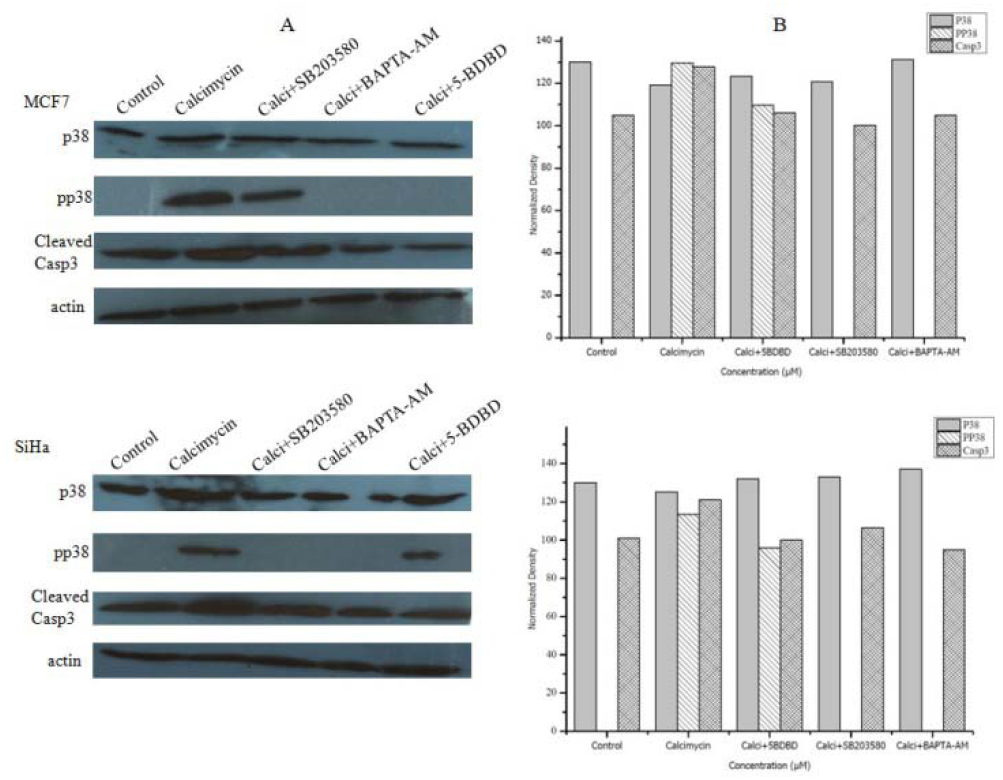
Western blot analysis of protein expression. (A) P38, phospho p38 (PP38), cleaved caspase 3 protein expression were determined in SiHa and MCF7 cell lines. β-actin is a loading control.(B) graph of band density values, normalized to actin signals

**Figure 15:**
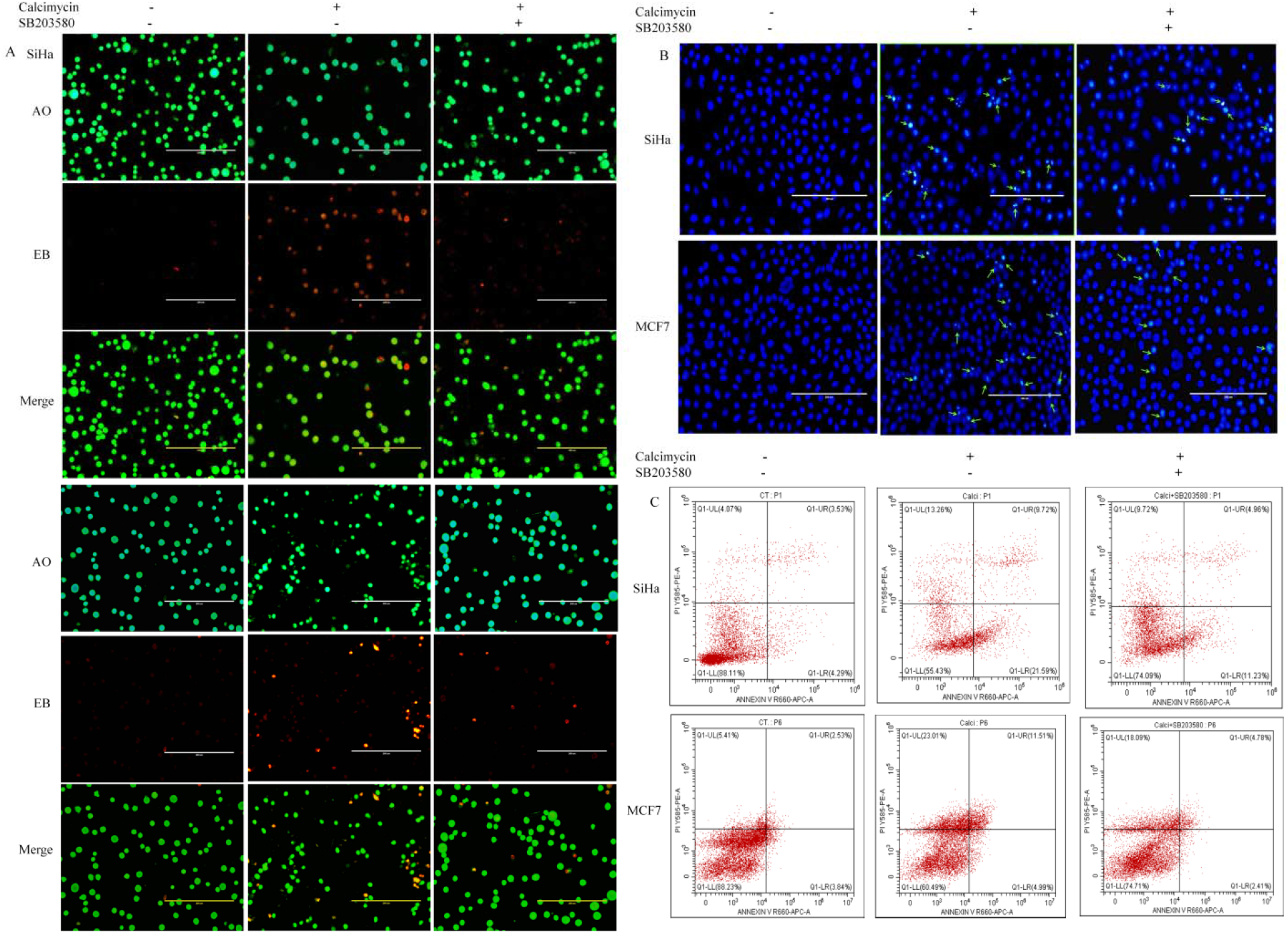
Effect of phospho P38 inhibitor SB203580 upon calcimycin induced apoptosis. (A) Effect of SB203580 on apoptosis by AO/EB staining. Early/late apoptotic cell no. get decreased in double positive cells i.e SB203580+calcimycin in comparison to calcimycin treatment only in both SiHa and MCF7 cell lines. (B) Effect of SB203580 on calcimycin mediated apoptosis by nuclear stain DAPI staining. In the effect of SB203580, cell shrinkage and chromatin fragmentation in SiHa and MCF7 get diminished as compared to calcimycin treatment only. Scale bar is 200 μm. (C) Flow cytometry assay shows late apoptotic cell no. get decreases in SB203580+calcimycin treated cells in comparison to only calcimycin treated cells

## 4. Discussion

This study set to determine the cytotoxic effects of calcimycin on human breast cancer cells and cervical cancer cells, and to find its mechanisms of action. On both malignant cell lines, cell viability analysis reveals that calcimycin inhibits cell proliferation in a dose-dependent manner. Calcimycin’s anti-metastatic potential on SiHa and MCF7 cell lines was also investigated using a cell migration inhibition in wound healing assay. Cell migration is exhibited by endothelial and epithelial monolayer movement in 2 dimensions that provide information about the rate of gap closure, which expedient the speed of collective motion of the cells (Jonkman et al., 2014). Calcimycin treatment was found to induce an extreme reduction in cell migration rate in both cell lines. These findings thus evidently demonstrate calcimycin’s antiproliferative potential in both cervical and breast cancer. We observed nuclear condensation, cytoplasm shrinkage, and apoptotic cell death in both cells in response to calcimycin using two different staining assays DAPI and AO/EB dual staining. Annexin V-APC/PI data substantiate the potential of calcimycin to exert apoptosis, especially in the late stage in both SiHa and MCF-7 cells as early apoptotic cells reduced on high concentration while late apoptotic cells increased. These data support the hypothesis of the cytotoxicity of calcimycin toward breast and cervical cancer through apoptosis in a dose-dependent manner. The cytosolic Ca^2+^ concentration is one of the main regulators of crucial cellular processes. Many physiological cell functions are monitored by Ca^2+^-mediated signaling. In cancer, Ca^2+^ signaling was found to be involved in several tumorigenesis events viz migration, angiogenesis, proliferation, and escape from apoptosis (Patergnani et al., 2020). Ca^2+^ signaling is one of the complex and interdependent cascades in cells whose elevation was proposed to be related to triggering apoptosis signals. Calcimycin is a calcium ionophore, therefore to determine the effect of calcimycin over intracellular Ca^2+^ levels, calcium measurement was done. Calcimycin treatment is found to induce significant increases in intracellular Ca^2+^ levels which get restored with prior treatment of calcium chelator BAPTA-AM. The further apoptotic assay also confirms the involvement of intracellular Ca^2+^ elevation in calcimycin induces apoptotic cell death as prior treatment of BAPTA-AM in both SiHa and MCF7 cells found to restore nuclear fragmentation, chromatin condensation and no. of late apoptotic cells. Ca^2+^ signaling is well known to be influenced and synchronized by mitochondria also. Mitochondrial membrane potential (Δψm), has a significant association with apoptosis, also assumed to be dominant in the apoptosis event. Depolarization of the mitochondrial membrane potential is induced by opening the transition pore within the mitochondrial membrane that releases apoptosis-inducing factors which makes a loss in Δψm as a consequence of apoptotic events (Ly et al., 2003). Δψm is intact in healthy cells where TMRM can pass to the mitochondria and accumulates in the inner mitochondrial membrane and gives high fluorescence intensity upon measurement by flow cytometry however in apoptotic or stressed cells TMRM gets distributed throughout the cytoplasm as it can no longer accumulate inside the mitochondria leading to less or reduced fluorescence compared to control (Verma & Das, 2018). Δψm is found to be reduced with increasing concentration of calcimycin in both cell lines pointing toward the involvement of mitochondrial consequence of apoptosis. In continuation with our experiments next, we try to explore the probable association of ATP receptor P2RX4 in calcimycin-mediated elevation of intracellular Ca^2+^. Several research studies demonstrated the linkage between ATP and intracellular Ca^2+^. ATP elevation has been reported to increase intracellular Ca^2+^ concentration in the cells through ligand-gated ion channels permeable to Ca^2+^ (Nishii et al., 2009). With reference to available reports, we presume to investigate the association of ATP release and involvement of ATP receptors in calcimycin-mediated intracellular Ca^2+^ elevation. For this we first checked the transcript level expression of P2RX4 in both cell lines in response to calcimycin treatment and found P2RX4 mRNA expression increased with increasing concentration of calcimycin treatment in both cell lines. These results led us to presume that calcimycin may enhance the ATP level as P2RX4 is a commonly known ATP-mediated receptor. To test this assumption we did ATP measurement and found significant enhancement in extracellular ATP level with increasing concentration of calcimycin in both cell lines. To bear out the possibility that calcimycin-mediated intracellular Ca^2+^ elevation responsible for apoptotic cell death is P2RX4 dependent ATP mediated pathway, we checked the intracellular Ca^2+^ level as well as apoptotic events with 1 hr prior treatment of P2RX4 antagonist 5-BDBD followed by 24 hrs incubation of calcimycin and found the deduction in intracellular Ca^2+^ as well as apoptotic events as compared to only calcimycin treated cells in both cell lines. These results suggest that calcimycin treatment raises extracellular ATP and intracellular Ca^2+^ levels also P2RX4 mRNA expression in both cancer cell lines, together with P2RX4 inhibition reduces intracellular Ca^2+^ level in cells plus calcimycin mediated apoptotic events, confirming the involvement of P2RX4 dependent ATP mediated pathway in calcimycin induced apoptotic cell death through raising intracellular Ca^2+^ in both cell lines. Molecular docking studies identify the highest and most stable binding potential between calcimycin and P2RX4 that controls ATP release to gain more understanding of how calcimycin regulates intracellular Ca^2+^ levels through P2RX4. These findings led us to the conclusion that calcimycin controls P2RX4 expression, increasing extracellular ATP levels and apoptosis flux. p38 MAPK has been considered to play a significant role in apoptotic signaling cascades. p38 MAPK has a considerable role in cell death as well as cell survival machinery. Depending on the type of stimulus as well as in a cell type-specific manner p38 MAPK can mediate cell survival and cell death too. Through the MAPK signaling pathway, Ca^2+^ up-regulation can stimulate a range of cellular responses; prompting either apoptosis or cell survival (Yue & López, 2020). These findings suggest the binary role of p38 which is specific to the external stimuli. To investigate the role of p38 MAPK activation in calcimycin-mediated apoptosis we checked the level of p38 & phosphorylated p38 (pp38) by western blotting. Western blot data shows calcimycin treatment increased the pp38 in both cell lines in comparison to untreated cells however no effect has been shown in the p38 level indicating that calcimycin increases the p38 phosphorylation. To confirm the p38 MAPK activation is upshots of calcimycin & related to intracellular Ca^2+^ increment & P2RX4 dependent ATP mediated pathway we pretreated the cells with BAPTA-AM and 5-BDBD as well as specific phosphorylated p38 inhibitor SB203580 and then treated with calcimycin for 24 hrs. Western blot analysis showing the attenuation of phosphorylated p38 as well as apoptotic marker cleaved caspase 3 in the presence of inhibitors & Ca^2+^ chelator with calcimycin as compared to only calcimycin treated cells. Both P2RX4 antagonist 5-BDBD and intracellular Ca^2+^ chelator attenuated the phosphorylated p38 expression level suggesting that the calcimycin causes activation of p38 phosphorylation by increasing intracellular Ca^2+^ level in P2RX4-dependent ATP mediated pathway. Next, to confirm the involvement of p38 phosphorylation in calcimycin-mediated apoptosis, other apoptotic assays like DAPI, AO/EB staining, and annexin V-APC/PI staining were also done with 2 hrs prior treatment of SB203580 followed by 24 hrs incubation with calcimycin and found Phosphorylated p38 specific inhibitor SB203580 attenuated the calcimycin induced apoptosis. Thereby confirming the apoptosis induced by calcimycin in both cancer cell lines are P2RX4 dependent ATP mediated intracellular Ca^2+^ and p38 MAPK mediated pathway. However, the precise mechanism by which calcimycin promotes p38 phosphorylation still needs further experimental confirmation.

## 5. Conclusion

The present study reveals the mechanism by which calcimycin induces apoptosis in breast and cervical cancer cell lines. The treatment of calcimycin increases the P2RX4 purinoreceptor mRNA expression as well as intracellular calcium level by mitochondrial Ca^2+^ influx in the cell which in turn activates phosphorylation of p38 and caspase 3 resulting in apoptotic cell death. Our results explored a new mode of action for calcimycin in cancer that could be potentially employed in future studies for cancer therapeutic research. This study disentangles that the calcimycin-induced apoptotic cell death or cell proliferation inhibition is P2RX4 dependent ATP mediated intracellular Ca^2+^ and p38 MAPK mediated pathway in both cell lines which is essentially concordant with the earlier study related to other cancer cell types. However, further research is needed to determine how the calcimycin affects p38 phosphorylation and P2RX4 activation.

## Acknowledgements

We acknowledge Central Discovery Centre (CDC), Banaras Hindu University, Varanasi for Flow Cytometry and Dr. Bhagyalaxmi Mohapatra, Associate Professor, Department of Zoology, Banaras Hindu University, Varanasi for Luminometer facility. We acknowledge Dr. Paresh Kulkarni, Department of Biochemistry, Institute of Medical Science, Banaras Hindu University for providing BAPTA-AM. We acknowledge Dr. Garima Jain, Malaviya Postdoctoral Fellow, Centre for Genetic Disorders, BHU for her critical reading and feedback on this manuscript.

## Compliance with ethical standards

### Author Contributions

Neha designed and performed the experiments, and wrote the manuscript. P.D. designed the experiment, review and supervision. P.R did the *In silico* experiments and wrote the manuscript.

## Appendix: List of Primers

P2RX4 FP: ATCCTTCCCAACATCACCAC P2RX4 RP: TGGCAAACCTGAAATTGTAGC

